# Structural assembly of maize CRY-GL2 photosignaling complex provides insights into its regulatory role in cuticular wax biosynthesis

**DOI:** 10.1101/2025.08.06.668860

**Authors:** Yaqi Liu, Zhiwei Zhao, Xue Zhang, Yahui Hao, Fan Feng, Yuan Chen, Ji Wang, Miaolian Ma, Jianxu Li, Fang Yu, Hongtao Liu, Peng Zhang

**Affiliations:** Key Laboratory of Plant Carbon Capture, CAS Center for Excellence in Molecular Plant Sciences, Institute of Plant Physiology and Ecology, Chinese Academy of Sciences, Shanghai, China; National Key Laboratory of Plant Molecular Genetics, Chinese Academy of Sciences, Shanghai, China; School of Life Science, Shenzhen University, Shenzhen, China; University of Chinese Academy of Sciences, Beijing, China; Shanghai Normal University, Shanghai, China; Shanghai Chenshan Plant Science Research Center, Chinese Academy of Sciences, Shanghai, China

**Keywords:** cryptochrome, photoreceptor, crystal structure, Glossy2, light signal, wax biosynthesis

## Abstract

Cryptochromes (CRYs) are blue-light photoreceptors in plants that regulate diverse physiological processes through oligomerization-dependent interactions with downstream effectors. While numerous light-inducible CRY-binding proteins have been identified, the structural basis for photoactivated CRY-effector assembly remains poorly understood. Here we report the crystal structure of a constitutively active maize CRY1c photolyase homology region (ZmCRY1c-PHR^W368A^) in complex with ZmGL2, a BAHD ptotein directing very-long-chain fatty acid (VLCFA) elongation for cuticular wax biosynthesis. Structural analysis reveals that light-activated ZmCRY1c forms a homotetrameric scaffold generating four symmetric binding sites at subunit interfaces. Each ZmCRY1c protomer engages one ZmGL2 molecule through conformational reorganization of its α16-α17 loop and α17 helix, establishing a 4:4 hetero-octameric photosignaling complex validated by mutational profiling and optical design. Remarkably, structural alignment with the ZmCER6-ZmGL2 complex reveals substantial overlap (78%) in ZmGL2 binding interfaces. Biochemical quantification demonstrates that ZmCRY1c suppresses ZmCER6-GL2 enzyme activity in a dose dependent manner, unveiling a light-dependent regulatory switch controlling VLCFA elongation efficiency, though *in vivo* physiological investigation is awaited. Collectively, our work not only establishes the atomic model of light-activated plant CRY-effector assembly, but also uncovers a previously unrecognized spatial competition between photoreceptor and metabolic enzyme complexes that might function as a photoregulatory paradigm in cuticular wax biosynthesis.

## Introduction

Cryptochromes (CRY) are photoreceptors that mediate blue light perception in plant and non-plant species(*1-3*). In plants, CRYs play major roles in hypocotyl elongation and floral initiation, as well as regulating stomatal opening and circadian rhythms(*4-7*). CRY proteins contain two domains, the photolyase homologous region (PHR) domain that noncovalently binds the chromophore FAD (flavin adenine dinucleotide) and the variable carboxy-terminal extension domain (CCE)(*8, 9*). In different plants, as well as Drosophila, zebrafish and mammals, the similarity of PHR domain is high, and the difference is mainly attributed to the CCE domain(*10, 11*).

Previous structural data together with the functional analyses have suggested the light activation process of CRY, in which the “trp-triad” residues mediate the blue-light dependent photoreduction of FAD, which induce conformational changes and oligomerization through the PHR domain, and finally the activation of CRY(*9, 12-15*). As an inhibitor of CRY, BIC could bind with CRY monomer and preclude the CRY oligomerization(*16-18*). The AtCRY2^W374A^ mutant, though photochemically inactive in photoreduction, could mimic the photoactivated conformation to form oligomers independent of light, and remain constitutively active(*14, 19*). The photoactivated or oligomerized CRY then bind with various downstream proteins to transmit light signal. To date, a large number of proteins have been identified to interact with plant CRYs in a light dependent manner(*20-29*). However, the molecular mechanisms underlying how photoactivated CRY interacts with downstream proteins to transmit light signals remain elusive due to the lack of atomic resolution structures between activated CRY and downstream protein complexes.

Here, we report the crystal structure of the maize CRY1c PHR domain in complex with a downstream effector GL2, a BAHD family protein required for the VLCFA elongation through forming a complex with ketoacyl-CoA synthase CER6 (*30-32*). The results obtained here made it clear on how the photoactivated CRYs to form a photsignaling complex and provides molecular insights on its regulatory role in adjusting the CER6-GL2 enzyme activity during VLCFA elongation and cuticular wax biosynthesis in maize.

## Results

### Active ZmCRY1c forms a complex with ZmGL2

It is remarkable to note that many CRY interaction proteins identified could not form a stable complex with CRY when recombinantly expressed in vitro for structural analysis, this brings difficulty to understand how CRY binds downstream effector to transmit photo signal. Luckily, a new hit Glossy2 (ZmGL2) appeared in search of CRY1 interaction proteins in maize, and ZmGL2 belongs to a BAHD acyltransferase superfamily member essential for VLCFA elongation and cuticular wax biosynthesis (*32-35*). Since mutation of residue Trp368 of ZmCRY1c to Ala (ZmCRY1c^W368A^) could enhance the oligomeric state which mimics the active conformation of ZmCRY1c(*14, 19*), we chose PHR domain of ZmCRY1c^W368A^ (ZmCRY1c-PHR^W368A^) for further analyses; and another FAD-binding failure mutant, ZmCRY1c-PHR^D381A^, was also generated based on sequence alignment with AtCRY2^D387A^ **(fig. S1)**. The above proteins were recombinantly expressed, and the photoreduction characteristics were measured (**Fig. 1A; and fig. S2A-C**). The results show that ZmCRY1c-PHR wild type features typical photoreduction, while ZmCRY1c-PHR^W368A^ and ZmCRY1c-PHR^D381A^ lost photoreduction under 50 μmol/m^2^/s blue-light condition. When recombinantly expressed, ZmGL2 form a stable complex with the PHR domain of the ZmCRY1c active mutant ZmCRY1c^W368A^ (ZmCRY1c-PHR^W368A^) in a 1:1 molar ratio (**Fig. 1B; and fig. S2D**). The binding affinity is of ZmCRY1c-PHR^W368A^ with ZmGL2 is about 0.6 μM (∼1.0 μM for ZmCRY1c^W368A^ full length), which is of several fold enhancement of wild type ZmCRY1c-PHR (KD∼2.8 μM; ∼2.7 μM for ZmCRY1c full length) (**Fig. 1C; and fig. S3**). Meanwhile, the FAD-binding failure mutant ZmCRY1c-PHR^D381A^, indicated by absence of photoreduction under 50 μmol/m^2^/s blue-light condition, lost the binding ability to ZmGL2 (**Fig. 1C**). All these data confirm the light dependent interaction between ZmGL2 and ZmCRY1c.

**Fig. 1.**
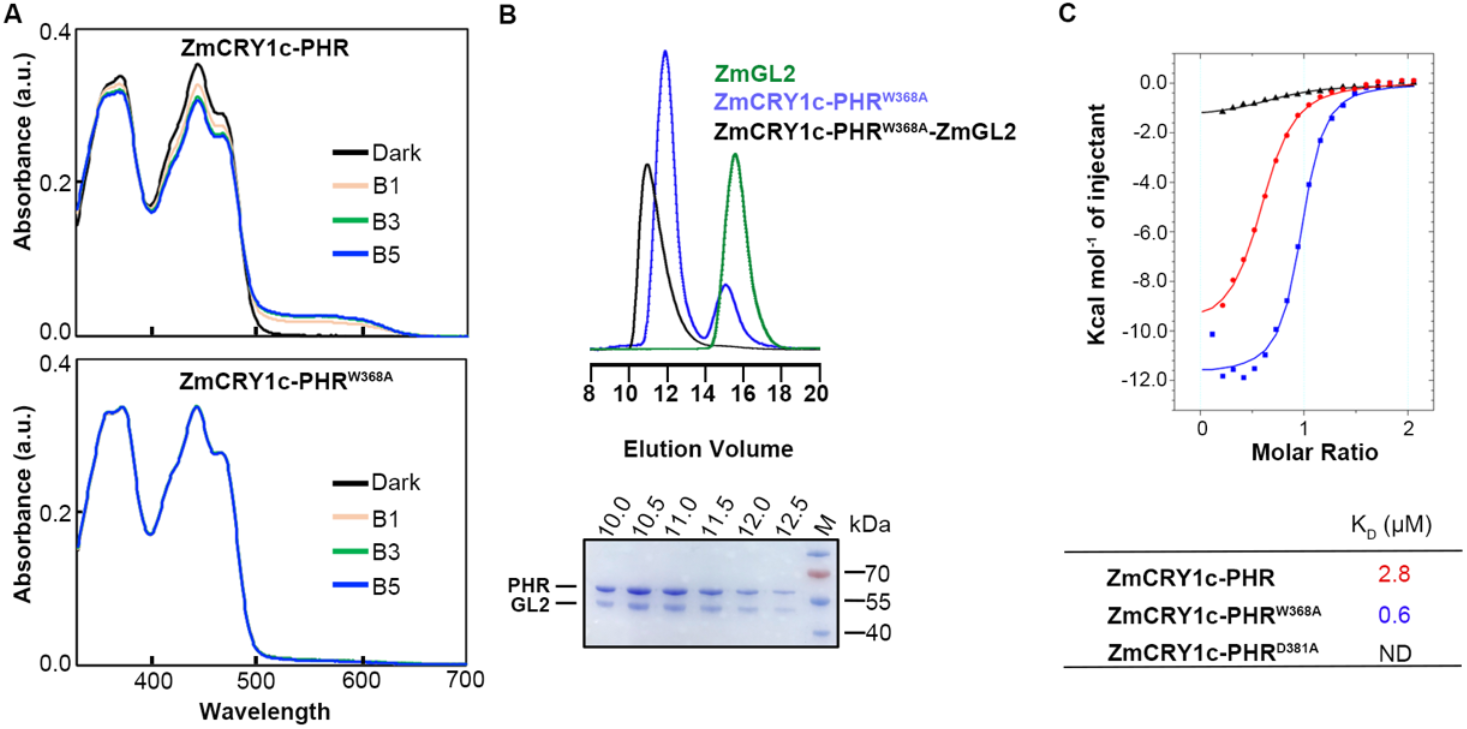
Reconstruction of the maize CRY1c-GL2 complex in *vitro*. **(A)** Photoreduction of wild type ZmCRY1c-PHR protein, ZmCRY1c-PHR^W368A^ protein. The absorption spectra of different protein were recorded at indicated times (0 min/Dark, 1 min/B1, 3 min/B3 and 5 min/B5) after blue-light illumination (50 μmol/m^2^/s^2^) under aerobic conditions at 20℃, 5 mM DTT was added as external electron donor. **(B)** ZmCRY1c-PHR^W368A^ interacted with ZmGL2 *in vitro*. SEC profiles of ZmCRY1c-PHR^W368A^-ZmGL2 complex (black), ZmCRY1c^W368A^-PHR (blue), ZmGL2 (green). Protein samples were eluted from Ni^2+^-chelating column applied to Superdex-200 10/300 for analysis (upper panels). Purified protein samples of ZmCRY1c-PHR^W368A^-ZmGL2 complex after SEC was resolved by SDS-PAGE and stained by Coomassie blue. Above SDS-PAGE gels, numbers indicate the peak fractions of the target protein (lower panels). **(C)** Binding affinity measurement of the ZmGL2 with ZmCRY1c-PHR (red curves), ZmCRY1c-PHR^W368A^ (blue curves), ZmCRY1c-PHR^D381A^ (black curves) using isothermal titration calorimetry, respectively. KD, dissociation constant. ND: Not detected. The syringe was filled with 500 μM ZmGL2; The cell was filled with 50 μM ZmCRY1c-PHR or their mutants.

### Architecture of ZmCRY1c-PHR^W368A^-ZmGL2 complex

The recombinant ZmCRY1c-PHR^W368A^-ZmGL2 protein complex was purified to homogeneity and concentrated to 10 mg/mL for crystallization. The crystal structure of ZmCRY1c-PHR^W368A^-ZmGL2 complex was determined to 2.8 Å resolution by molecular replacement method **(Fig. 2A-F**, and **fig. S4, A and B)**. Based on the obtained structure information, we found that ZmCRY1c-PHR^W368A^ forms a homologous tetramer through crystallographic stacking **(Fig. 2A-C)**. As expected, the ZmCRY1c-PHR^W368A^ tetrameric crystal structure is overall similar to the ZmCRY1c-PHR cryo-EM structure (PDB: 6LZ3) previously determined in active conformation (RMSDs of 1853 Cα atoms are 1.19 Å**) (Fig. 2D)**. Two interaction interfaces INT1 (A/B or C/D) and INT2 (A/C or B/D) between ZmCRY1c-PHR monomers defined before(*14*) were used here for consistency. INT1 is constituted by helices α6-α7, α12-α13, α15, α18-α20 from monomers A/B or C/D, while INT2 is constituted by helices α2, α10, α5-α6 connecting loop from monomers A/C or B/D (**fig. S5, A and B**). ZmGL2 adopts a typical BAHD conformation (**fig. S4B**) and binds symmetrically near the INT1 of ZmCRY1c-PHR^W368A^ (**Fig. 2, E and F**), which forms a ZmCRY1c-PHR^W368A^-ZmGL2 hetero-octameric complex with 4:4 molar ratio.

**Fig. 2.**
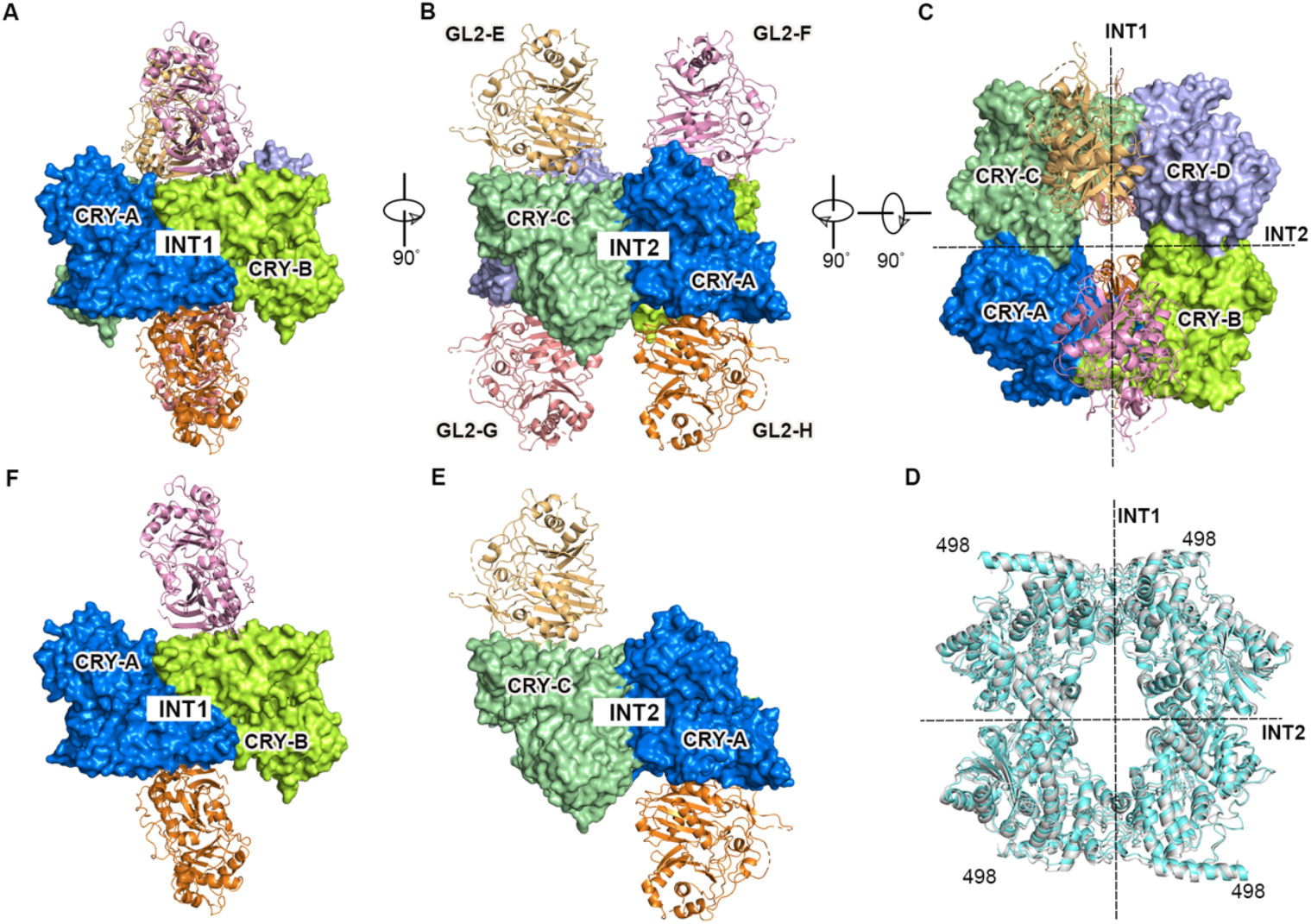
Architecture of ZmCRY1c-PHR^W368A^-ZmGL2 complex. **(A)-(C)** Hetero-octamer structure of ZmCRY1c-PHR^W368A^-ZmGL2 complex in different views. Front view **(A)**, side view **(B)** and top view **(C)**. The four CRY molecules A, B, C, D are colored by marine, limon, pale green and light blue, respectively. The four ZmGL2 molecules E, F, G, H are colored by light-orange, pink, salmon, and orange respectively. ZmCRY1c-PHR^W368A^ molecules and ZmGL2 molecules are shown with surface and ribbon, respectively. INT1: interface 1; INT2: interface 2. The INT1 and INT2 dimer interfaces are indicated with dotted lines. The drawing is done with pymol. **(D)** Superimposition of ZmCRY1c-PHR^W368A^ tetramer crystal structure (cyan) with the ZmCRY1c-PHR^W368A^ cryo-EM tetramer structure (6LZ3, gray). **(E)-(F)**Hetero-tetramer structure of ZmCRY1c-PHR^W368A^-ZmGL2 complex.

### Light dependent specific interaction between maize CRY1c and GL2

The interaction interface between ZmCRY1c-PHR^W368A^ and ZmGL2 mainly involves α16-α17 loop, α17 of ZmCRY1c (residues 396-419), and α2, β1-β2 loop of ZmGL2, which buries 640 Å^2^ of solvent accessible surface area **(Fig. 3, A and B)**. The interactions between ZmCRY1c-PHR^W368A^ and ZmGL2 are mainly hydrophobic, and are composed of residues Arg406, Ile407, Asn409, Gln411, Phe412, Tyr415 from ZmCRY1c-PHR^W368A^ and residues Val25, Ser27, Val29, Phe71 from ZmGL2 **(Fig. 3B)**. Comparing the crystal structure of ZmCRY1c-PHR^W368A^ in the ZmCRY1c-PHR^W368A^-ZmGL2 complex with the ZmCRY1c-PHR^W368A^ cryo-EM structure (PDB: 6LZ3), we found that the α17 and α16-α17 loop of ZmCRY1c undergo conformational changes due to ZmGL2 binding **(Fig. 3C)**. Specifically, the side chains of residues Phe412, Gln411 and Tyr415 of ZmCRY1c flip significantly to form interactions with residues of ZmGL2. This is consistent with our previous data suggested that CIB1 binding leads to conformational change of active AtCRY2(*36*). It is worth noting that the interaction-participating structural elements of ZmCRY1c bridge the α16 on which the “trp-triad” residue Trp391 located with the α18-α20 that make up the INT1 **(Fig. 3D)**. This may explain why the activated ZmCRY1c induces tight binding of ZmGL2.

**Fig. 3.**
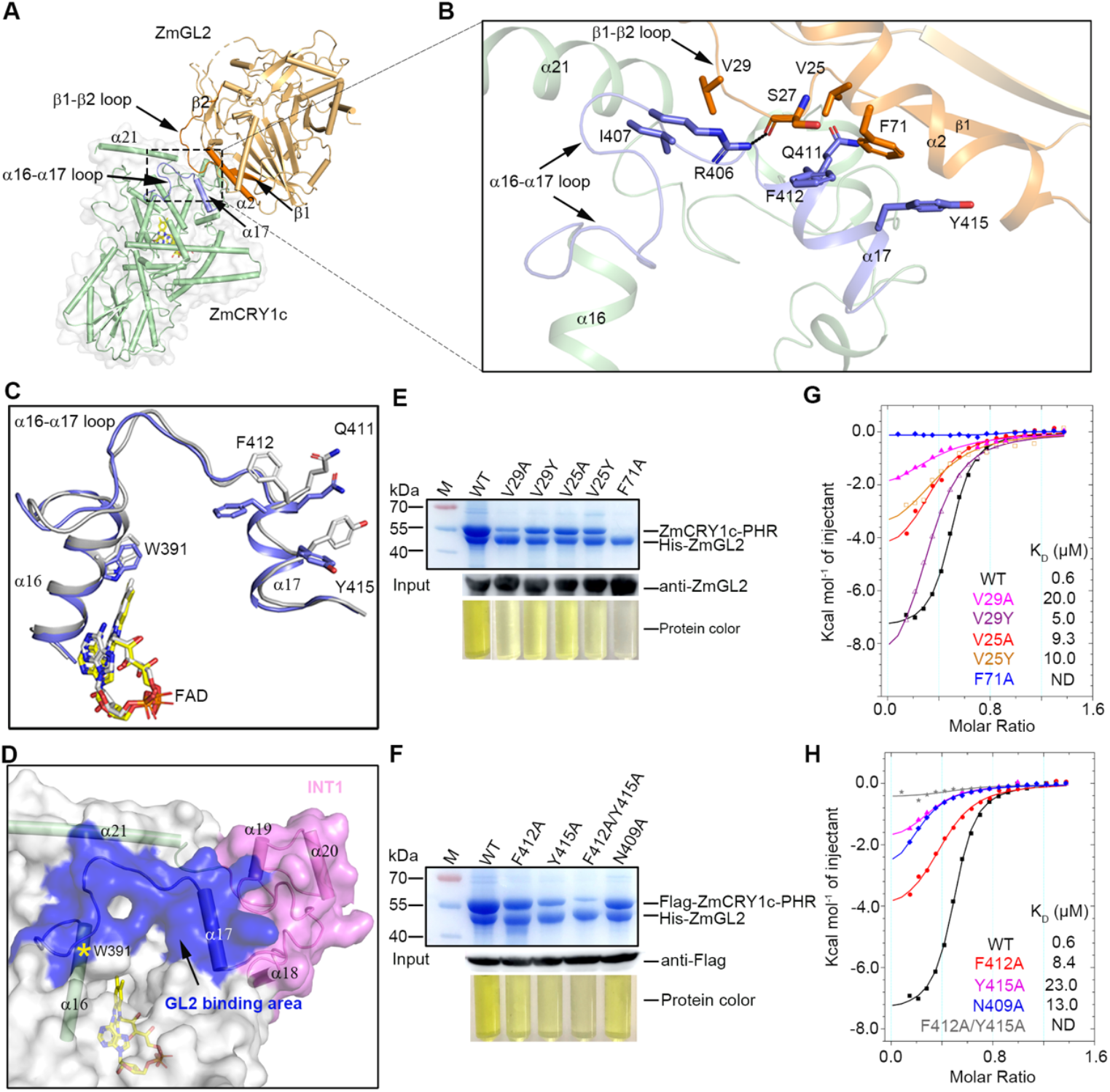
Structure and biochemistry analysis of CRY-GL2 interface. **(A)-(B)** ZmCRY1c-PHR^W368A^ and ZmGL2 interaction interface. **(A)** ZmCRY1c-PHR^W368A^ are shown with surface and cylindrical helices. ZmGL2 are shown with cylindrical helices. ZmCRY1c-PHR^W368A^ and ZmGL2 are colored with palegreen and light orange, respectively. Structural elements constituting the interaction interface, α16-α17 loop and α17 from ZmCRY1c-PHR^W368A^ are colored with slate blue, and β1, β1−β2 loop, β2 and α2 from ZmGL2 are colored with orange. **(B)** A zoom-in view of ZmCRY1c-PHR^W368A^ and ZmGL2 interaction residues. Residues involving interaction are highlighted by side chains, and colored slate and orange, respectively. **(C)** A close-up view of the conformational changes at the α16-α17. ZmCRY1c-PHR^W368A^ crystal structure and Cryo-EM structure were colored with blue and gray. **(D)** A close-up view of the ZmGL2 binding site (blue) near the INT1 (magenta) of ZmCRY1c, the position of Trp391 is denoted as a yellow star. **(E)** His pull-down analysis on the ZmCRY1c-PHR^W368A^ and ZmGL2 interaction. SDS--PAGE gel shows the influence of His-ZmGL2 site mutations on the complex formation (upper panel). The protein color of the resulting complex is shown in the lower panel. The input of ZmGL2 protein and mutants is indicated using an anti-ZmGL2 antibody (middle panel). **(F)** His pull-down analysis on the ZmCRY1c-PHR^W368A^ and ZmGL2 interaction. SDS-PAGE gel shows the influence of Flag-ZmCRY1c-PHR^W368A^ (WT for short) site mutations on the complex formation (upper panel). The protein color of the resulting complex is shown in the lower panel. The input of Flag-ZmCRY1c-PHR protein and mutants is indicated using an anti-Flag antibody (middle panel). **(G)-(H)** Binding affinity measurement of the mutants referred in **(E)** and **(F)** using ITC, respectively. KD, dissociation constant. ND: Not detected. The syringe is filled with 1 mM ZmGL2 or their mutants; The cell is filled with 150 μM ZmCRY1c-PHR or their mutants.

To verify whether the binding is specific, we mutated the interaction residues and detected their influence on the interaction between ZmCRY1c-PHR^W368A^ and ZmGL2 using His pull-down analysis **(Fig. 3, E and F)**. The results showed that mutation of residue Val 29 or Phe 71 from ZmGL2 to Ala could severely impair or destroy the interaction between ZmCRY1c-PHR^W368A^ and ZmGL2, while less effects were found in the other mutations V29Y, V25A or V25Y **(Fig. 3E)**. Meanwhile, double mutation of residues Phe412 and Tyr415 from ZmCRY1c to Ala could also severely impair the interaction, while single mutation Y415A, F412A, or N409A has less effect on the interaction **(Fig. 3F)**. The mutational effects were also indicated by the faded yellow color of ZmCRY1c protein. To quantitatively assess the influence of these mutations on ZmCRY1c-PHR^W368A^-ZmGL2 complex formation, the binding affinity in between was measured using ITC experiments (**Fig. 3, G and H**). As can be seen from the results, the binding affinity between ZmCRY1c-PHR^W368A^ and the mutant ZmGL2^V29A^ or ZmGL2^V25Y^ or ZmGL2^V25A^ or ZmGL2^V29Y^ was decreased by ∼33, ∼17, ∼16, ∼8 folds, respectively; and the ZmGL2^F71A^ mutant even lost binding capacity with ZmCRY1c-PHR^W368A^ **(Fig. 3G)**. Meanwhile, the binding affinity between ZmGL2 and ZmCRY1c-PHR^W368A^ harboring mutation Y415A, N409A, or F412A was decreased by ∼33, ∼22, ∼14 folds, respectively, and the double mutation F412A/Y415A may lost binding with ZmGL2 **(Fig. 3H)**. All these data confirm that the interactions observed in the ZmCRY1c-PHR^W368A^-ZmGL2 crystal structure are specific.

The specific interaction between ZmCRY1c and ZmGL2 was further tested and used for an optical design. A simplified system was adopted in which ZmCRY1c-PHR and ZmGL2 genes were fused respectively to BD and AD that control the LacZ expression in yeast under blue light(*37, 38*) **(Fig. 4A)**. The growing yeast cells were treated by blue light or dark for time slots, and the β-galactosidase activity was determined which reflects the complex formation between ZmCRY1c-PHR and ZmGL2. The results suggest that in contrast to the wild type which activates the LacZ expression to a high level in the presence of 30 μmol/m^2^/s blue light, the activation ability of ZmGL2^V25A^ mutation was reduced to ∼50 precent, while the ZmGL2^V29A^ and ZmGL2^F71A^ mutations showed no activation ability (**Fig.4B**). Meanwhile, the activation ability of ZmCRY1c^F412A^ and ZmCRY1c^F412A/Y415A^ were reduced respectively to ∼42 and ∼12 precent of wild type, while the ZmCRY1c^D381A^ mutation lost activation ability (**Fig. 4C**). The background activation in the dark was due to the difficulty in avoiding light during experiment process (**fig. S6**). These data not only confirm the light dependent interaction between ZmCRY1c and ZmGL2, but also provide the concept of their usage in optical design.

**Fig. 4.**
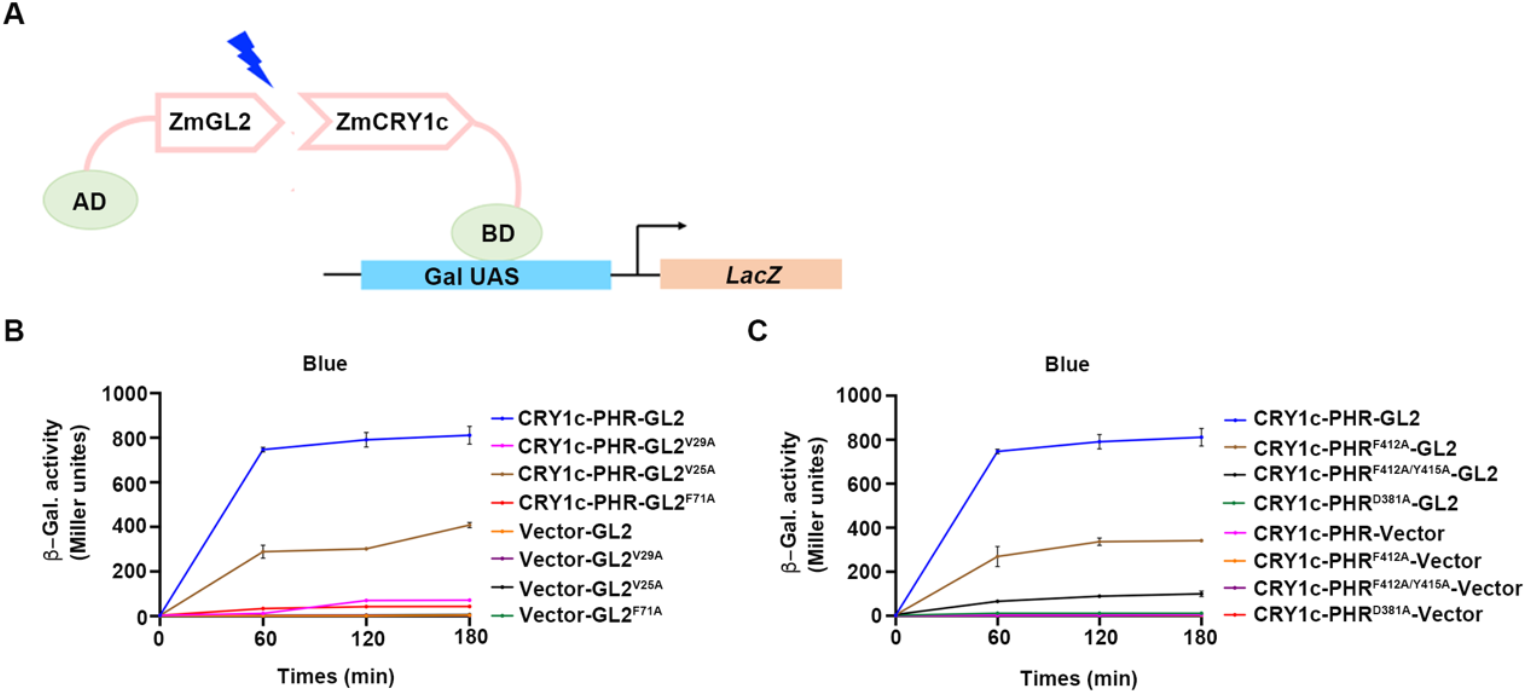
Optical switch design using ZmCRY1c and ZmGL2. **(A)** Schematic optical switch model of ZmCRY1c and ZmGL2. ZmCRY1c-PHR and ZmGL2 genes are fused respectively to BD and AD that control the *LacZ* expression in yeast under blue light. **(B)-(C)** β-galactosidase activity of different constructs.

Next, we want to test whether the interactions between CRY and GL2 exist in other species. However, we found that the interaction residues are not conserved among plant CRYs or GL2-like proteins (**figs. S7 and S8**). To further verify this observation, we purified the *Arabidopsis* constitutive active mutant AtCRY2^W374A^ and tested its interaction with the ZmGL2 homologue in *Arabidopsis* AtCER2 *in vitro* using ITC experiment. As expected, no interaction could be detected between AtCRY2-PHR^W374A^ and AtCER2, or AtCRY2-PHR^W374A^ and ZmGL2, or ZmCRY1c-PHR^W368A^ and AtCER2 (**fig. S9**). Surprisingly, we found that a chimera in which the 401-489 amino acids of AtCRY2^W374A^-PHR was replaced with 395-500 amino acids of ZmCRY1C^W368A^-PHR (containing the ZmGL2 binding site and INT1 region that covers the α16-α17 loop and helices α17-α21) could interact with ZmGL2, and the dissociation constant was determined to be ∼31 μM. This suggests that the interaction between CRY and GL2 may specifically exist in maize.

### ZmCRY1c adjusts ZmCER6-ZmGL2 enzyme activity through competing with ZmCER6

ZmGL2 has been established as a central regulator of wax biosynthesis by mediating VLCFA elongation via interaction with ketoacyl-CoA synthase CER6 (*32, 33, 35*). Our parallel cryo-EM study reveals the ZmCER6-GL2 heterotetramer architecture, demonstrating how ZmGL2 binds ZmCER6 to expand the substrate-binding tunnel of ZmCER6 to accommodate >C28 acyl-chains and enable VLCFA elongation. Remarkably, structural alignment revealed that ZmCRY1c and ZmCER6 share 78% interface overlap on ZmGL2’s N-terminal domain (**Fig. 5, A-C**), suggesting potential competitive binding. Since ZmCER6-GL2 complex integrity is essential for enzymatic activity, we proposed that light-activated ZmCRY1c regulate VLCFA elongation through competitive binding with ZmCER6.

**Fig. 5.**
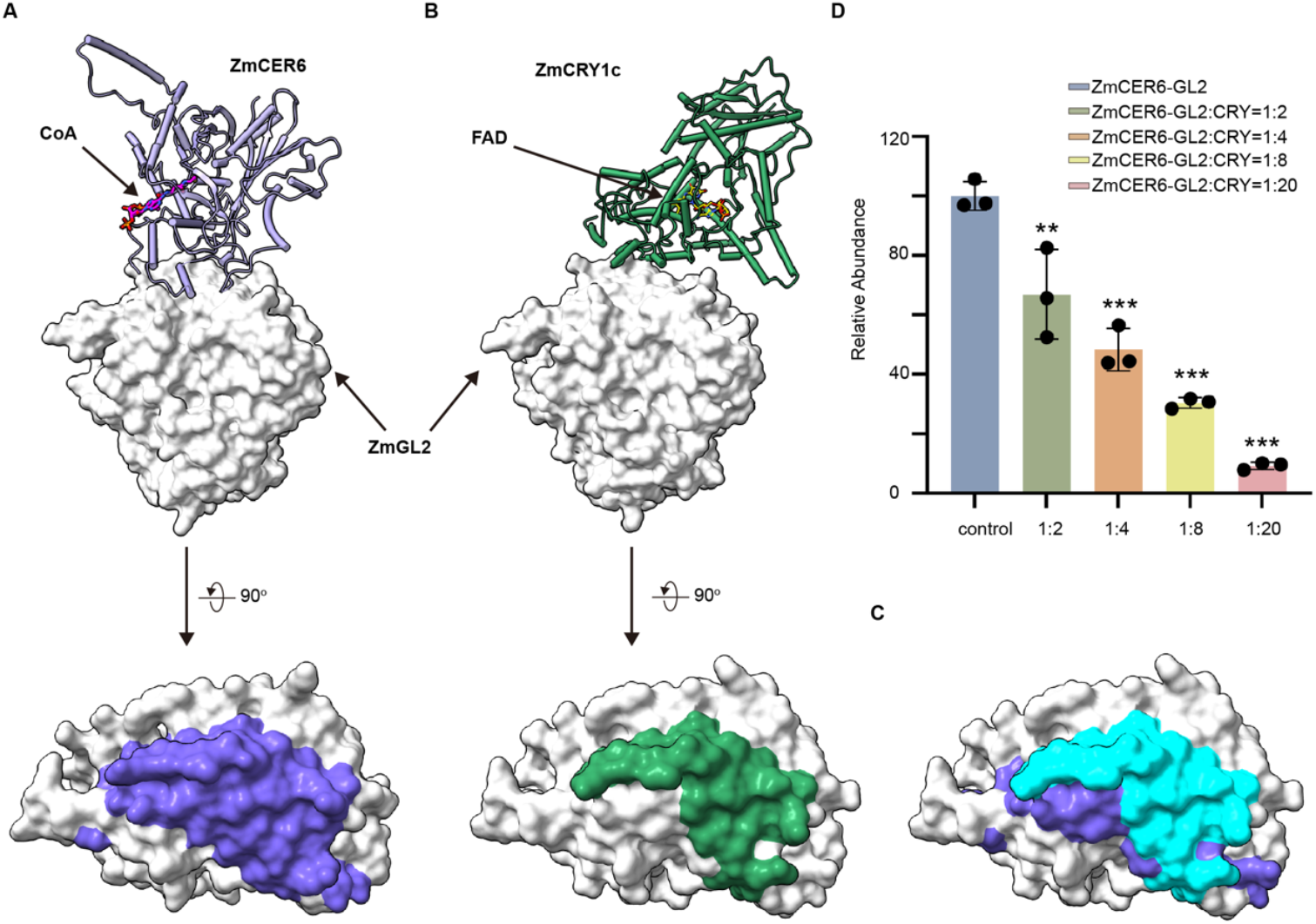
ZmCRY1c adjusts ZmCER6-ZmGL2 activity through competing with ZmCER6. **(A)** Binding site of ZmCER6 on ZmGL2. The ZmCER6-ZmGL2 complex structure is shown (upper panel); ZmCER6 is shown as a ribbon cartoon colored slate-blue, while ZmGL2 is shown as a surface model colored gray. The binding site of ZmCER6 on ZmGL2 is indicated with a blue surface patch (lower panel). **(B)** Binding site of ZmCRY1c on ZmGL2. The ZmCRY1c^W368A^-PHR-ZmGL2 complex structure is depicted with ZmGL2 adopting a similar orientation to that observed in the ZmCER6-ZmGL2 complex (upper panel). ZmCRY1c is shown as a ribbon cartoon colored green. The binding site of ZmCRY1c on ZmGL2 is indicated with a green surface patch (lower panel). **(C)** The overlapping binding interface between ZmCER6 and ZmCRY1c on ZmGL2 is shown as a cyan surface patch. **(D)** Inhibitory effect of ZmCRY1c on the ZmCER6-GL2 enzyme activity. 32:0-ketoacyl-CoA (m/z [M+H]+ 1244.5818, product, RT: 3.8 min) was detected using LC-MS assays. Different amount of ZmCRY1c (ratios are ZmCRY1c to ZmCER6) was added into the enzyme assay system.

Enzymatic assays demonstrated a dose-dependent inhibition of ZmCER6-GL2 activity by ZmCRY1c, with >90% suppression achieved at 20:1 molar ratio (**Fig. 5D**). This competitive inhibition mechanism suggests light-activated ZmCRY1c acts as a molecular switch displacing ZmCER6 from ZmGL2, thereby modulating VLCFA elongation reaction. While *in vivo* plant validation studies are awaited, these findings suggest that ZmCRY1c may function as a light-dependent switch in regulating cuticular wax biosynthesis through adjusting the VLCFA elongation efficiency.

## Discussion

While over thirty CRY-interacting proteins have been identified through genetic and biochemical studies(*26, 39, 40*), the structural basis for photoactivated CRY-effector complex assembly has remained poorly characterized. Our work bridges this critical gap by establish the atomic-resolution structure of a plant cryptochrome bound to its downstream effector ZmGL2, a BAHD protein forming complex with the ketoacyl-CoA synthase ZmCER6 to exert condensation activity during VLCFA elongation and cuticular wax biosynthesis. Moreover, we establish that light-activated ZmCRY1c directly competes with the ZmCER6 for ZmGL2 binding, creating a dose-dependent regulatory circuit that gates VLCFA elongation efficiency, a biochemical mechanism reconciling CRY’s dual roles in photoperception and metabolic regulation.

Structural and biochemical analyses demonstrate that ZmCRY1c oligomerization generates a ZmGL2-binding interface involving the α16-α17 loop and α17 helix proximal to the INT1 region **(Fig. 3, A-D)**, confirming our prior prediction of INT1 as a functional interaction nexus (*14*). This interface comprises residues evolutionarily conserved in plant CRYs but absent in animal homologs, implying a lineage-specific signaling adaptation. Intriguingly, these elements spatially align with the C-terminal lid of *Drosophila* CRY (dCRY) (*41-43*), a region mediating monomeric dCRY-TIM interactions in circadian regulation (**fig. S10**), and share topological parallels with photolyase DNA-binding modules (*44*), suggesting an evolutionary conservation of this structural module across different photoactivated CRYs. The mechanistic convergence extends to *Arabidopsis* CRY2, where blue light-induced oligomerization exposes the INT2 surface for CIB1 effector recruitment (*36*). Collectively, these observations support a paradigm wherein photoactivated CRY oligomerization dynamically generates distinct interaction surfaces (INT1/INT2), enabling context-specific recruitment of diverse effectors-a mechanistic rationale for CRY’s pleiotropic signaling roles.

The ZmCRY1c-ZmGL2 structure also delineates a spatial linkage between the ZmGL2-binding interface and the conserved tryptophan triad (W315/W368/W391) forming an electron-transfer chain from the FAD pocket to INT1 **(fig. S11A)**. We propose a stepwise activation mechanism: blue light triggers flavin semi-reduction (FADH•), inducing conformational changes that propagate through the Trp triad to dissociate the CCE domain from PHR. Subsequent INT1 restructuring drives CRY1 oligomerization, repositioning the α16-α17 structural elements to establish the ZmGL2-binding interface (**fig. S11B**). GL2 binding then stabilizes this active conformation through residue-level adjustments, completing the photocycle.

This mechanistic model acquires biological significance through GL2’s dual partnerships. ZmGL2 binds ZmCER6 to expand the substrate-binding tunnel of ZmCER6 by creating a continuous hydrophobic channel at the ZmCER6-GL2 interface, thereby enable its accommodation of >C28 acyl-chains and VLCFA elongation, a prerequisite for cuticular wax deposition. The overlapping ZmGL2-binding surfaces of ZmCRY1c and ZmCER6, coupled with ZmCRY1c’s competitive inhibition of ZmCER6-GL2 activity, position the ZmCRY1c-GL2 axis as a photoregulatory node controlling wax biosynthesis flux. While *in planta* validation is further required, our findings suggest a paradigm wherein light quality directly modulates cuticular wax biosynthesis through CRY-mediated metabolic complex competition, an optoregulatory mechanism potentially critical for plant environmental adaptation.

## Materials and methods

### Gene cloning and Protein purification

The genes encoding ZmCRY1c, ZmGL2 were cloned from maize B73 cDNA by PCR. For protein expression in *E. coli*, genes were subcloned into pET-duet vector, with a 6×His tag at the N terminus of ZmGL2. The point mutation was generated by one-step PCR. The AtCRY2-ZmCRY1c chimera was generated by overlap PCR. *E. coli* BL21 (DE3) strain was used for protein expression. The bacteria were cultured at 37℃, 200 rpm and induced by 0.25 mM isopropyl β-D-thiogalactopyranoside (IPTG) for 12 h. The bacteria were collected and resuspended in buffer A (20 mM Tris-HCl, pH 8.0, 100 mM NaCl, 1 mM TCEP) supplemented with 1 mM PMSF, and lysed by a high-pressure cell disruptor at 700 bar and centrifuged at 20,000 g for 1 h. The supernatant was incubated with Ni-NTA beads (Qiagen) for 1 h in 4℃. After that, Ni-NTA beads with the bound protein was washed by buffer A supplemented with 25 mM imidazole. The protein was eluted by buffer A supplemented with 250 mM imidazole and concentrated for further purification on a Superdex-200 10/300 column.

### Crystallization, data collection and structure determination

The purified ZmCRY1c-PHR^W368A^-ZmGL2 complex was concentrated to 10 mg/mL for crystallization. Initial protein crystals were obtained at 20℃ in condition of 20% (w/v) Polyethylene glycol 3,350, 100 mM Bis-tris Propane/Hydrochloric acid pH 6.5, 200 mM Sodium fluoride. The protein to reservoir ratios is 1:1 in three-well sitting-drop crystallization plates. For crystal optimization, 1.5 μL protein mix with 1.5 μL reservoir was set up in 24-well hanging drops plates. The crystals were cryoprotected by reservoir solutions supplemented with 30% glycerol, and flash-cooled in liquid nitrogen. Data was collected at BL19U1 beamline of the Shanghai Synchrotron Radiation Facility (SSRF) under 100 K liquid nitrogen stream. The data were processed with HKL3000 package(*45*), and the initial phase was determined by molecular replacement with Phenix, using the cryo-EM structure of ZmCRY1c^W368A^ (PDB ID: 6LZ3) and AlphaFold(*46*) predicted structure of ZmGL2 (AlphaFold ID: AF-A0A804MAL4-F1) as templates. The ZmCRY1c-PHR^W368A^-ZmGL2 complex structure model was manually built with Coot(*47*). The complex structure was refined by manual adjustment in coot and refinement with Phenix(*48*). Data collection and refinement statistics are reported in Extended Data Table1.

### Isothermal titration calorimetry

Isothermal titration calorimetry (ITC) was performed with Microcal ITC 200 (Malvern) at 20℃. The cell of machine was filled with 200 μL ZmCRY1c-PHR and their mutants, respectively. The syringe was filled with 40 μL ZmGL2 and their mutants, respectively. All samples were centrifuged at 14000 g for 10 min before titration. The titration was performed by injections of 2 μL aliquots followed by 120 s of equilibration; the reaction was conducted 20 times injections. The reaction heat change was measured after individually injection. The data was treated and analyzed by Origin.

### UV-Vis Absorption Spectroscopy

UV-Vis absorption spectra of the recombinantly expressed protein ZmCRY1c-PHR, ZmCRY1c-PHR^W368A^, and ZmCRY1c-PHR^D381A^ were measured with UV-Vis spectrophotometer (UV-2600, Shimadzu, Japan). 600 μL protein samples (6 mg/ml) in buffer containing 20 mM Tris-HCl, pH 8.0, 100 mM NaCl, 5 mM DTT were kept at quartz cuvettes with an optical path length of 1 cm. Samples were illuminated using LED lamps with wavelengths of 450∼470 nm and power of 8.3 W to irradiate protein samples. The vertical distance between the light source and the sample is ∼24 cm. The absorption spectra (300 nm-700 nm) of samples were recorded at indicated times (0 min, 1 min, 3 min and 5 min) after blue-light illumination (50 μmol/m^2^/s) under aerobic conditions at 20°C. The absorption curves were plotted using GraphPad.

### Yeast two-hybrid assay

The complementary DNA (cDNA) fragments of ZmGL2 and mutants were cloned into the pGADT7 vector, and the fragments of ZmCRY1c-PHR and mutants were cloned into the pBridge vector. Recombinant vectors were co-transformed into yeast Y190 (*Saccharomyces cerevisiae*) strain and plated on the Trp-Leu dropout solid medium. The plates were incubated at 30℃ for 3-4 days. Yeast colonies were then selected and suspended on Trp-Leu dropout liquid medium supplement with 0.02% adenine and 2% glucose. Selected colonies were grown in dark conditions overnight before transfer to YPD media. Cells in YPD media was divided into dark and blue light groups, then cultured in 28℃, 200 rpm for around 2 hours until OD600 obsorption reached 0.5. β-galactosidase activity was measured using chlorophenol red-β-D-galactopyranoside (CPRG) as substrate. Three biological replicates were used for each combination.

### *In vitro* enzyme activity assays

The *in vitro* enzymatic assays for inhibitory effect of ZmCER6-GL2 complex by ZmCRY1c^W368A^-PHR protein were performed in a 50 µL total volume. The reaction system containing 40 μM 30:0-CoA, 40 μM malonyl-CoA substrates, 10 µM purified ZmCER6-GL2 protein, ZmCRY1c^W368A^-PHR protein (20 µM, 40 µM, 80 µM and 200 µM protein was added, respectively) and 100 mM Tris-HCl buffer (pH 7.5).

## Data availability

The atomic coordinates of ZmCRY1c-PHR^W368A^-ZmGL2 complex structure have been deposited in the Protein Data Bank with accession code 9LPG.

## Acknowledgements

We thank the staff members at BL18U1/BL19U1-SSRF for their technical assistance in X-ray diffraction data collection. We also thank professor B. Ding, Z. Chen and S, Tang at the school of Chemistry and Chemical Engineering of the Shang Hai Jiao Tong University. This work was supported by grants from the National Key R&D Program of China (grant no. 2024YFA1306701 to F.Y.), the National Natural Science Foundation of China (grant nos. 32230050 and 32025020 to P.Z.), the Chinese Academy of Sciences (grant no. XDB27020103 to P.Z.), and the Shanghai Science and Technology Commission (grant no. 23310710100 to P.Z.).

## Author Contributions

Y.L. designed the experiments and carried out the bulk of the experiments; Z.Z. and F.F. contributed to data analysis; Y.H., Y.C. and J.W. contributed to protein expression, purification, and optical design experiments; X.Z. and J.L. contributed structure determination; M.M. contributed to ITC experiments. P.Z. H.L. and F.Y analyzed the data and wrote the manuscript with inputs from other authors. P.Z. conceived the project.

**The authors declare no competing interests**.

