## Supplemental information for "Structural assembly of maize CRY-GL2 photosignaling complex provides insights into its regulatory role in cuticular wax biosynthesis"

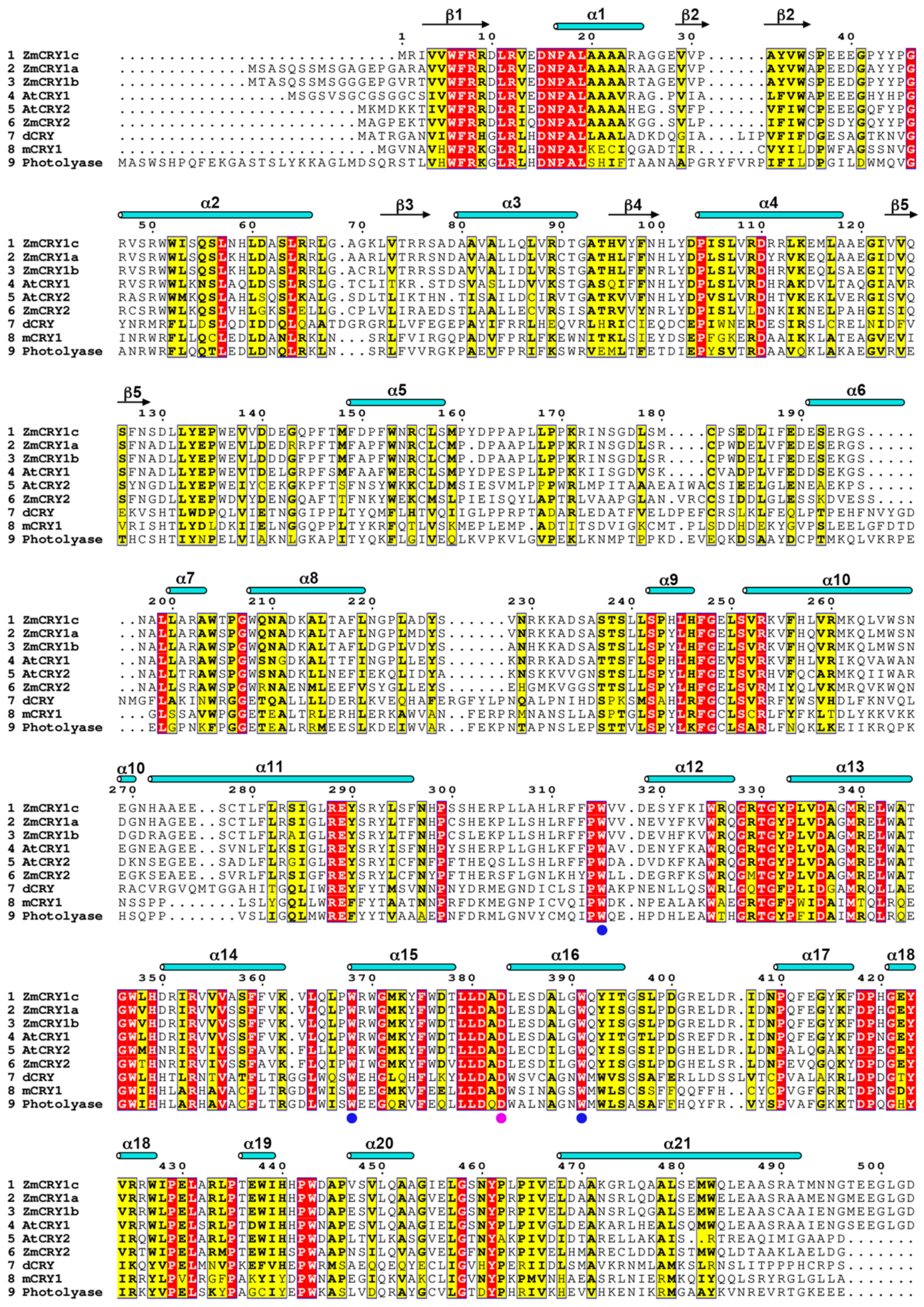


**Fig. S1. Sequence alignment of the PHR domain of cryptochromes and homologs from different species.**

Invariant residues and conserved residues are shaded with red box and yellow box, respectively. Trp-triad residues (Trp315, Trp368, Trp391) are indicated with blue circles, and residues (Asp381) play key roles in FAD binding is indicated with magenta circles. Secondary structure of ZmCRY1c is shown on the top.

Zm, *Zea mays*, ZmCRY1c (Uniprot: B8A2L5), ZmCRY1a (Uniprot: A0A1D6HB66), ZmCRY1b (Uniprot: K7TXG5), ZmCRY2 (Uniprot: K7VU84); At, *Arabidopsis thaliana*, AtCRY1 (Uniprot: Q43125), AtCRY2 (Uniprot: Q96524); dCRY (Uniprot: O77059), *Drosophila melanogaster* CRY; mCRY1 (Uniprot: P97784), mouse CRY; Photolyase (Uniprot: Q0E8P0), *Drosophila melanogaster* Photolyase.

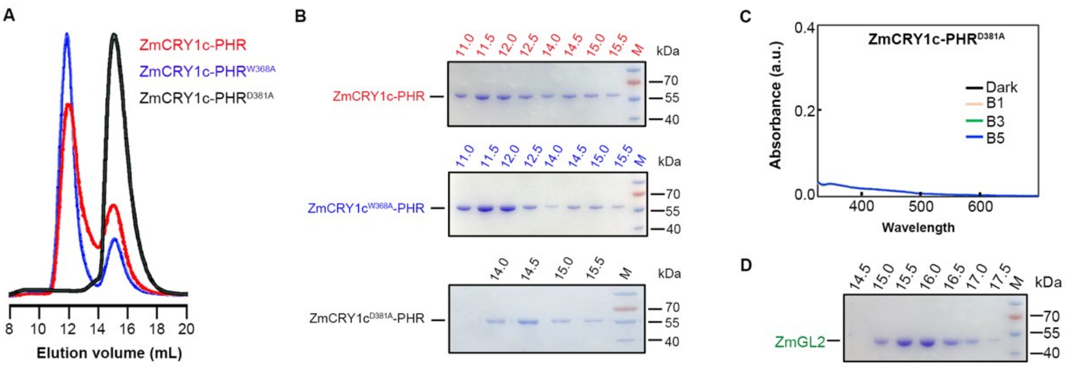


**Fig. S2. Purification and photoreduction of ZmCRY1c.**

**(A)** Size-exclusion chromatography (SEC) profiles of ZmCRY1c-PHR (red), constitutive active mutant ZmCRY1c-PHR$^{W368A}$ (blue), constitutive inactive ZmCRY1c-PHR$^{D381A}$ (black).

**(B)** Purified protein samples after SEC were resolved by SDS-PAGE and stained by Coomassie blue. Above SDS PAGE gels, numbers indicate the peak fractions of the target protein.

**(C)** Photoreduction of ZmCRY1c-PHR$^{D381A}$ protein. The absorption spectra of protein were recorded at indicated times (0 min, 1 min, 3 min and 5 min) after blue-light illumination (50 μmol/m$^2$/s$^2$) under aerobic conditions at 20°C, 5 mM DTT was added as external electron donor.

**(D)** Purified ZmGL2 protein samples after SEC were resolved by SDS-PAGE and stained by Coomassie blue, numbers above indicate the peak fractions of SEC.

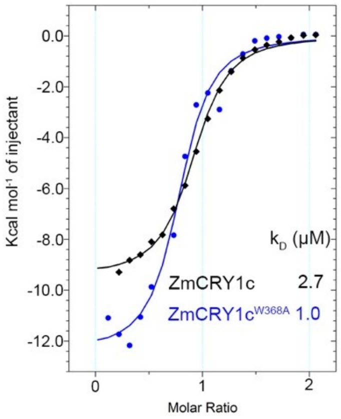


**Fig. S3. Binding affinity measurement of ZmGL2 with ZmCRY1c and ZmCRY1c$^{W368A}$.**

ZmCRY1c and ZmCRY1c$^{W368A}$ were colored by black and blue, respectively. $K_D$, dissociation constant.

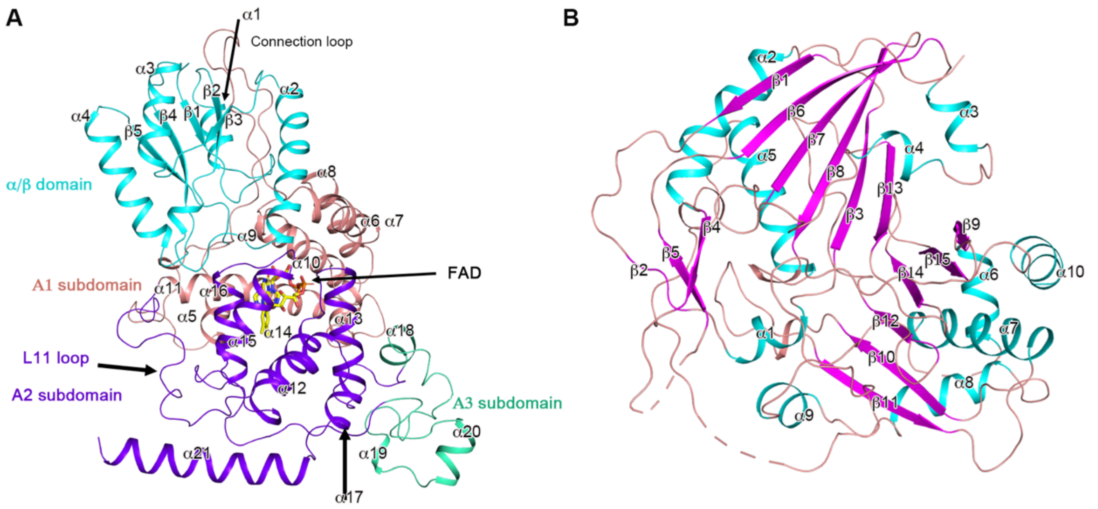


**Fig. S4. Monomer structure of ZmCRY1c-PHR$^{W368A}$ and ZmGL2.**

**(A)** Monomer structure of ZmCRY1c-PHR$^{W368A}$. The α/β domain and the A1, A2, and A3 subdomains of ZmCRY1c$^{W368A}$-PHR are colored as cyan, salmon, purple-blue, green, respectively.

**(B)** Monomer structure of ZmGL2. Helix, sheet, and loop were colored cyan, magenta, and salmon, respectively.

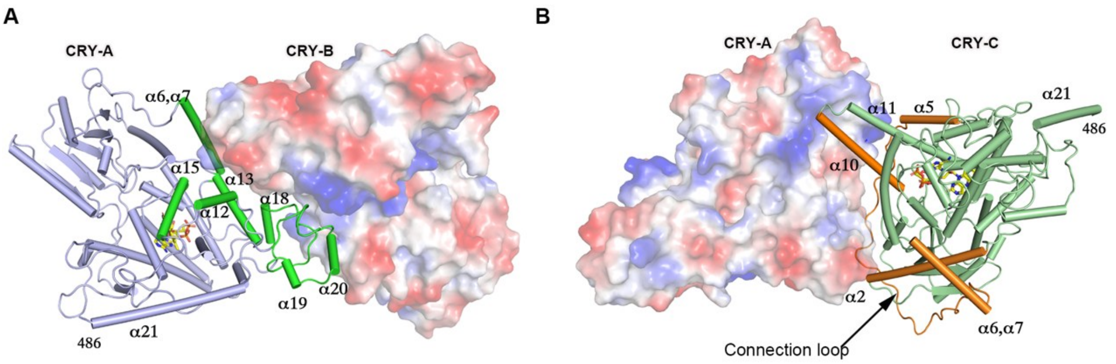


**Fig. S5. Interaction surface I and II (INT1/2) in ZmCRY1c-PHR$^{W368A}$ tetramer.**

**(A)** INT1 of ZmCRY1c-PHR$^{W368A}$ between monomers A and B. Monomer A is show as cylindrical helices (blue) and the interaction region is colored green; Monomer B is show as surface model.

**(B)** INT2 of ZmCRY1c-PHR$^{W368A}$ between monomers A and C. Monomer A is shown as surface model; Monomer C is shown as cylindrical helices (green) and the interaction region is colored orange.

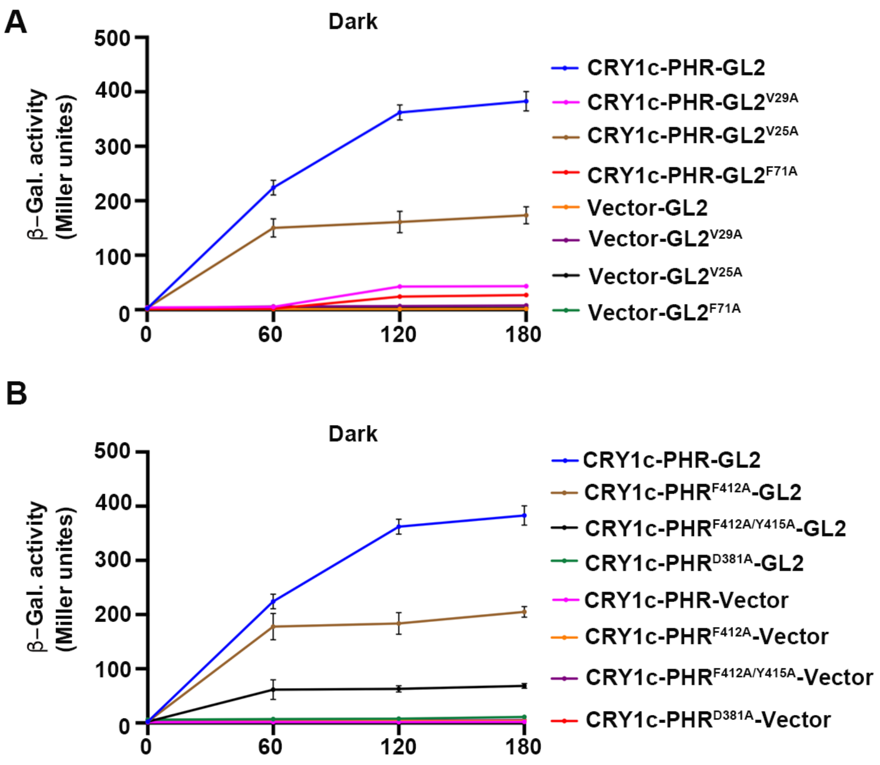


**Fig. S6. β-galactosidase activity of different constructs in dark.**

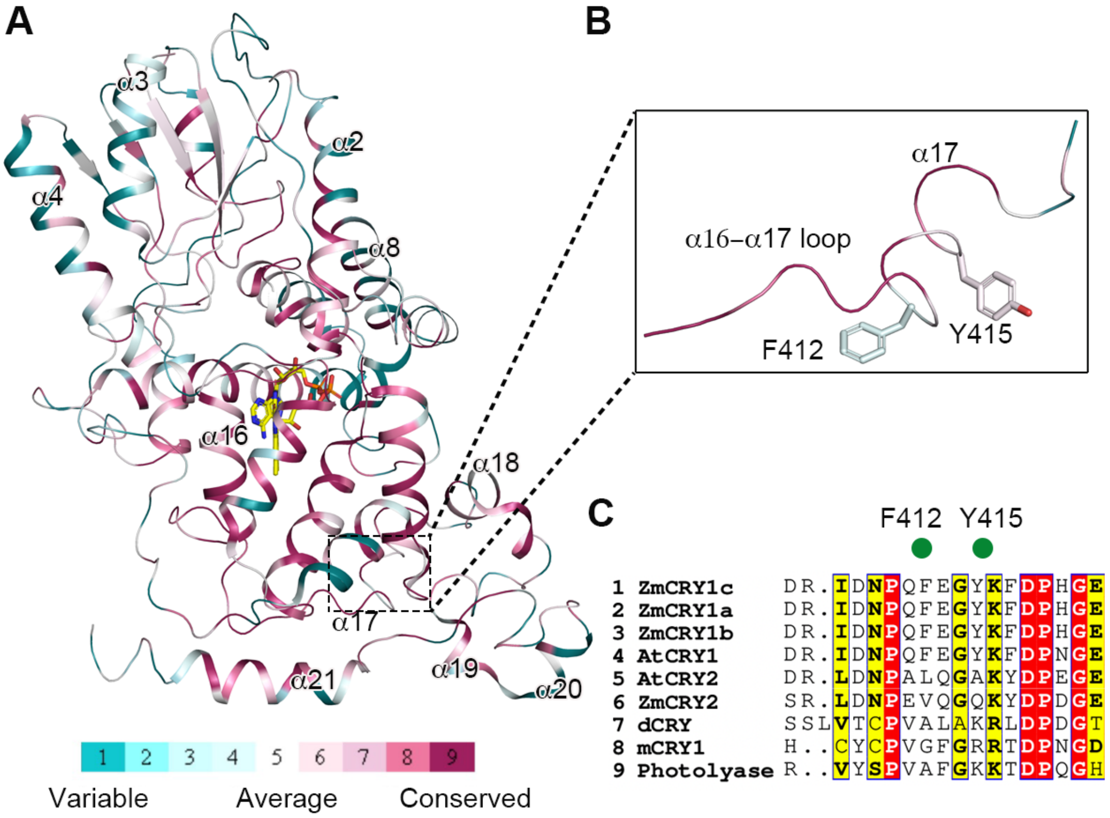


**Fig. S7. Sequence conservation analysis of the PHR domain of CRY from different species.**

**(A)** CRY PHR domain was colored by the conservation scores of amino acids using ConSurf web server.

**(B)** A zoom-in view of the side chains of F412 and Y415, which are located at α17 of ZmCRY1c.

**(C)** Sequence alignment of α16-α17 loop and α17 from different species. Amino acids sequences alignment using CLUSTALW and ESPript 3.0 online software, High conserved residues are red background, while less conserved residues are red colored, not conserved residues are black colored. Zm: *Zea mays* L., At: *Arabidopsis thaliana*. Os: *Oryza sativa* L.

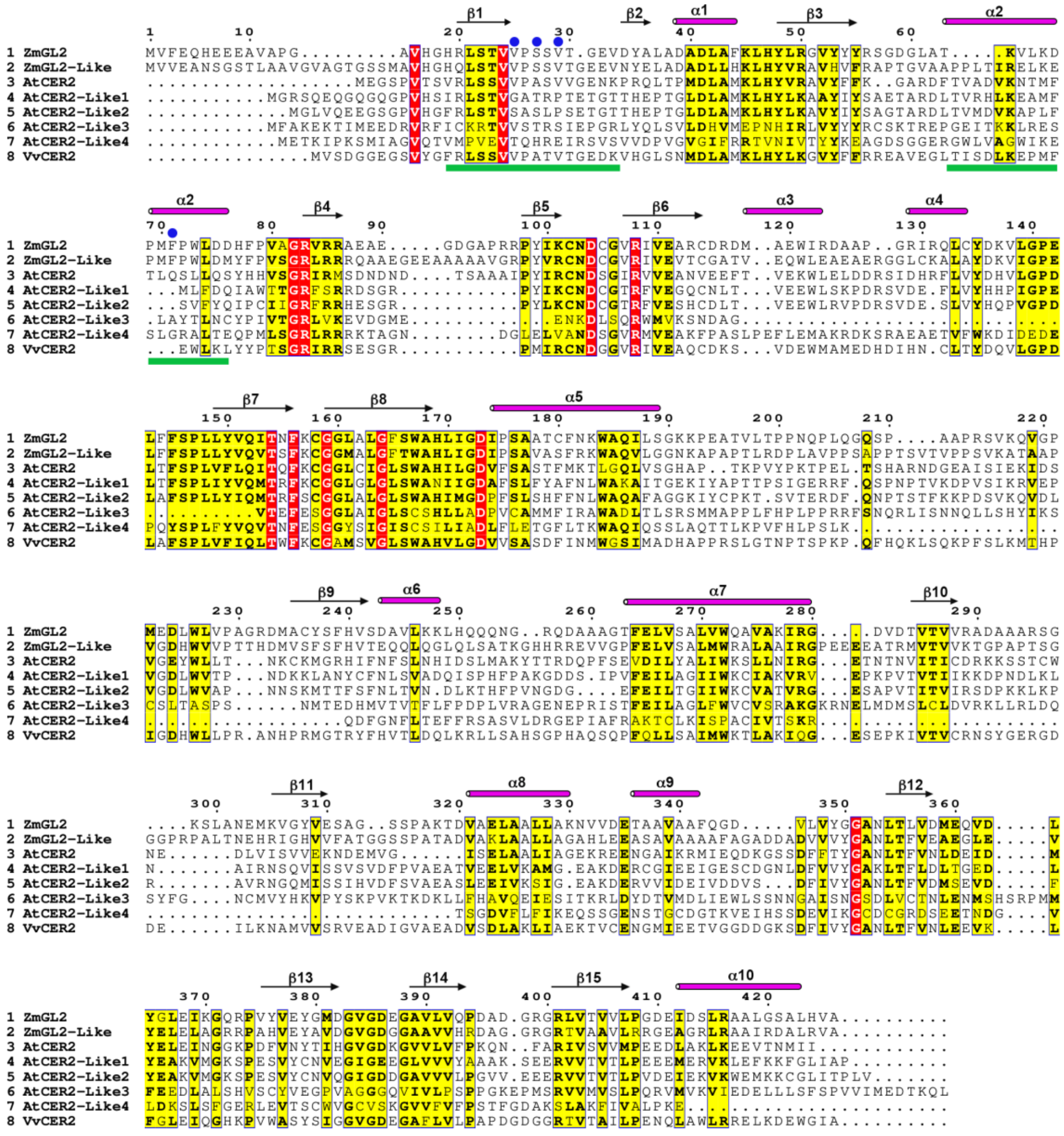

**Fig. S8. Sequence alignment of ZmGL2 protein and homologs from different species.**

Invariant residues and conserved residues are shaded with red box and yellow box, respectively. Structure elements and key residues involving in interacting with ZmCRY1c are indicated with green rectangles and blue circles, respectively. Zm, *Zea mays*, ZmGL2 (Uniprot: A0A804MAL4), ZmGL2-Like (Uniprot: A0A804RBN5); At, *Arabidopsis thaliana*, AtCER2 (Uniprot: Q39048), AtCER2-Like1 (Uniprot: Q9SVM9), AtCER2-Like2 (Uniprot: Q9LIS1), AtCER2-Like3 (Uniprot: F4JMX1), AtCER2-Like4 (Uniprot: Q9LYZ6); Vv, *Vitis vinifera*, VvCER2 (Uniprot: E0CRX2).

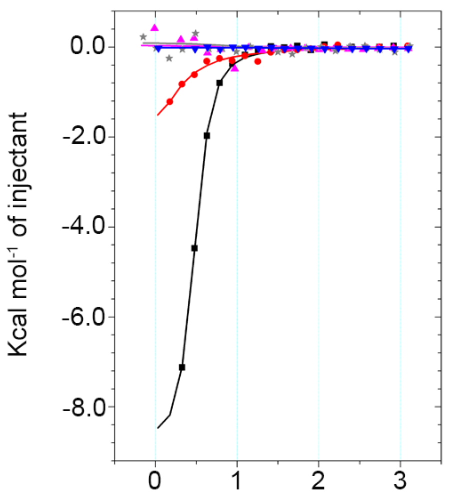


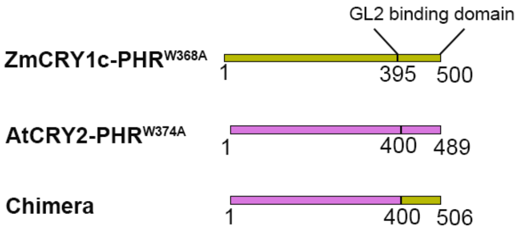


| Mol-A | Mol-B | $K_D$ (μM) |
|---|---|---|
| AtCRY2-PHR$^{W374A}$ | AtCER2 | ND |
| AtCRY2-PHR$^{W374A}$ | ZmGL2 | ND |
| ZmCRY1c-PHR$^{W368A}$ | AtCER2 | ND |
| ZmCRY1c-PHR$^{W368A}$ | ZmGL2 | 0.6 |
| Chimera | ZmGL2 | 31.0 |

**Fig. S9. Binding affinity measurement of different protein constructs.**

The binding curves of AtCRY2-PHR$^{W374A}$ and AtCER2 (blue)/ ZmGL2 (gray), ZmCRY1c-PHR$^{W368A}$ and AtCER2 (magenta)/ZmGL2 (black), AtCRY2$^{W374A}$ (1-400)-ZmCRY1c (395-500) (chimera) and ZmGL2 (red), respectively (left panel). Construct of chimera is shown in the top right panel. $K_D$, dissociation constant. ND: Not detected. The syringe is filled with 1.5 mM ZmGL2 or their mutants, and the cell is filled with 100 μM ZmCRY1c-PHR or their mutants.

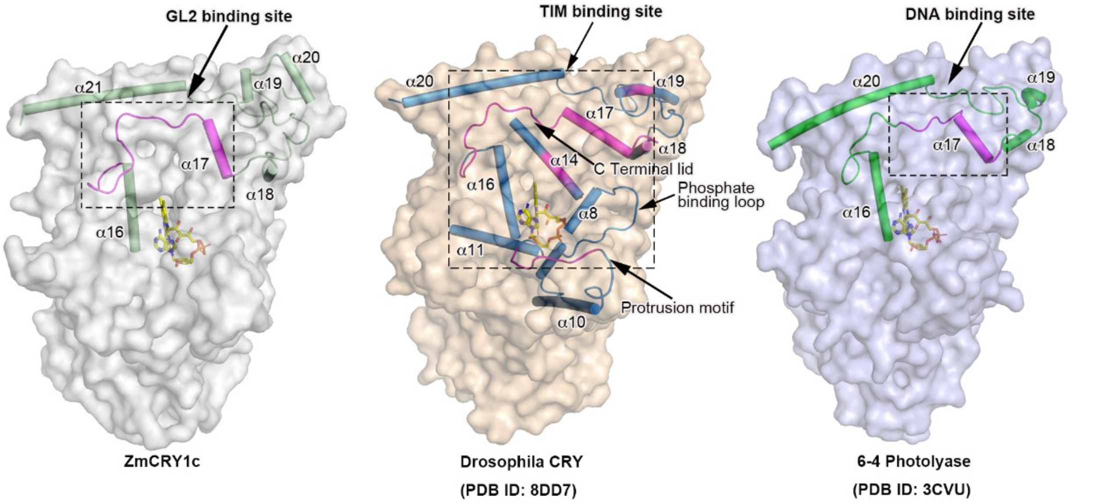


**Fig. S10. Different CRYs or Photolyase bind downstream effector proteins or DNA using a similar region**

ZmCRY1c (left panel), dCRY (middle panel) and 6-4 Photolyase (right panel) were show with surface models at similar orientation. The binding sites of ZmGL2, Timeless (TIM) and DNA are indicated and colored magenta.

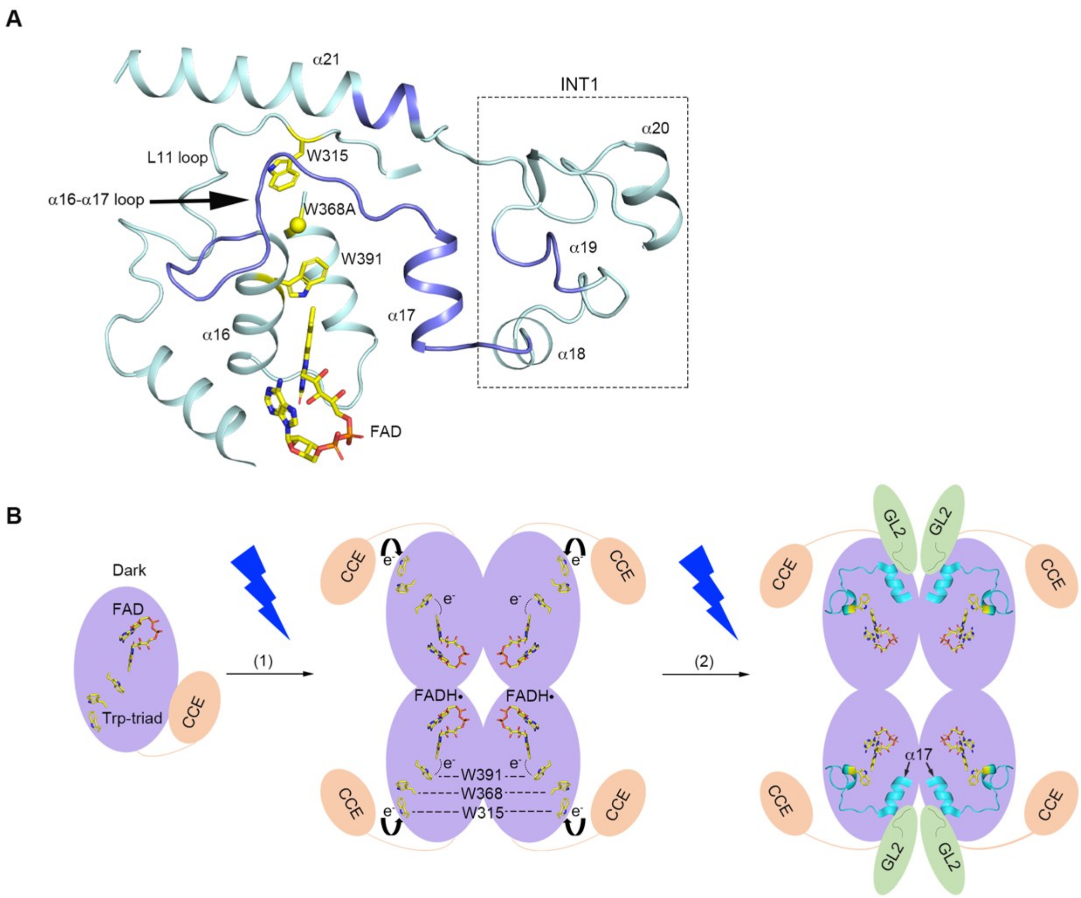


**Fig. S11. Photoactivated ZmCRY1c interacts with ZmGL2 to transmit photo signal**

**(A)** A close-up view of the FAD and "trp-triad" residues (W391/W368A/W315) relative location with α16-α17 loop and α17 helix.

**(B)** A proposed model of ZmCRY1c-ZmGL2 photosignaling complex transmit signal.

### Table S1. Data collection and model refinement statistics

| | ZmCRY1c-PHR-ZmGL2 |
|---|---|
| **Data collection** | |
| Space group | $C222_1$ |
| Cell dimensions | |
| a,b,c (Å) | 108.34, 217.05, 228.80 |
| α,β,γ (°) | 90, 90, 90 |
| Resolution (Å) | 42.17-2.80 (2.90-2.80) [a] |
| Rmerge | 0.308 (1.429) |
| I/σ (I) | 8.33 (1.33) |
| $CC_{1/2}$ | 0.975 (0.641) |
| Completeness (%) | 91.50 (49.74) |
| Redundancy | 13.0 (10.7) |
| **Refinement** | |
| Resolution (Å) | 42.17-2.80 (2.90-2.80) |
| No. reflections | 60946 |
| $R_{work}$ / $R_{free}$ | 0.2317/0.2675 |
| No. atoms | 13981 |
| Protein residues | 1761 |
| Ligands | 106 |
| Water | 33 |
| r.m.s. deviations | |
| Bond lengths (Å) | 0.007 |
| Bond angles (°) | 1.17 |
| Average B-factor | 66.93 |
| Protein | 67.28 |
| Ligands | 32.83 |
| Water | 31.21 |
| Ramachandran | |
| Favored (%) | 94.14 |
| Allowed (%) | 5.86 |

[a] Values in parentheses are for highest-resolution shell
